# SARS-CoV-2 transmission via apical syncytia release from primary bronchial epithelia and infectivity restriction in children epithelia

**DOI:** 10.1101/2021.05.28.446159

**Authors:** Guillaume Beucher, Marie-Lise Blondot, Alexis Celle, Noémie Pied, Patricia Recordon-Pinson, Pauline Esteves, Muriel Faure, Mathieu Métifiot, Sabrina Lacomme, Denis Dacheaux, Derrick Robinson, Gernot Längst, Fabien Beaufils, Marie-Edith Lafon, Patrick Berger, Marc Landry, Denis Malvy, Thomas Trian, Marie-Line Andreola, Harald Wodrich

## Abstract

The beta-coronavirus SARS-CoV-2 is at the origin of a persistent worldwide pandemic. SARS-CoV-2 infections initiate in the bronchi of the upper respiratory tract and are able to disseminate to the lower respiratory tract eventually causing acute severe respiratory syndrome with a high degree of mortality in the elderly. Here we use reconstituted primary bronchial epithelia from adult and children donors to follow the infection dynamic following infection with SARS-CoV-2. We show that in bronchial epithelia derived from adult donors, infections initiate in multi-ciliated cells. Then, infection rapidly spread within 24-48h throughout the whole epithelia. Within 3-4 days, large apical syncytia form between multi-ciliated cells and basal cells, which dissipate into the apical lumen. We show that these syncytia are a significant source of the released infectious dose. In stark contrast to these findings, bronchial epithelia reconstituted from children donors are intrinsically more resistant to virus infection and show active restriction of virus spread. This restriction is paired with accelerated release of IFN compared to adult donors. Taken together our findings reveal apical syncytia formation as an underappreciated source of infectious virus for either local dissemination or release into the environment. Furthermore, we provide direct evidence that children bronchial epithelia are more resistant to infection with SARS-CoV-2 providing experimental support for epidemiological observations that SARS-CoV-2 cases’ fatality is linked to age.

**Significance Statement:** Bronchial epithelia are the primary target for SARS-CoV-2 infections. Our work uses reconstituted bronchial epithelia from adults and children. We show that infection of adult epithelia with SARS-CoV-2 is rapid and results in the synchronized release of large clusters of infected cells and syncytia into the apical lumen contributing to the released infectious virus dose. Infection of children derived bronchial epithelia revealed an intrinsic resistance to infection and virus spread, probably as a result of a faster onset of interferon secretion. Thus, our data provide direct evidence for the epidemiological observation that children are less susceptible to SARS-CoV-2.

## Introduction

Coronaviruses with zoonotic origin have emerged as a new public health concern during the first decades of the 21th century. Two highly pathogenic coronaviruses, severe acute respiratory syndrome coronavirus (SARS-CoV) and Middle-East respiratory syndrome coronavirus (MERS-CoV), caused severe respiratory infections in humans during regionally confined epidemics in 2002 (1) and between 2010-15 (2), respectively. In late 2019, clusters of patients with pneumonia in Wuhan in the Hubei province in China were shown to be infected with the novel severe acute respiratory syndrome coronavirus 2 (SARS-CoV-2) (3–5). SARS-CoV-2 infections are associated with acute respiratory illness referred to as <u>Co</u>rona<u>vi</u>rus <u>d</u>isease (COVID-19). Since its description, SARS-CoV-2 infections are at the root of an enduring worldwide pandemic, having caused as of May 2021 over 3 million deaths and more than 148 million confirmed infections (data from the John Hopkins university coronavirus resource center, https://coronavirus.jhu.edu/). SARS-CoV-2 is an enveloped virus with a positive single-stranded RNA of around 30 kb. The 5’ proximal two thirds of the polyadenylated genome encodes ORF1a and ORF1b, which are autoproteolytically processed into several non-structural proteins required for replication and transcription. The distal third encodes for the 4 structural proteins, Envelope (E), Membrane (M), Nucleocapsid (N) and Spike (S) and seven putative ORFs encoding accessory proteins and potential virulence factors (6–8). The surface exposed Spike protein gives the virus its crown-like appearance in electron microscopy and mediates the attachment to the main cellular receptor ACE2 (9). Coronaviruses can cause a wide range of respiratory illnesses, from mild upper respiratory tract infection up to a severe acute respiratory syndrome (10). The latter is characterized by excessive cytological damage and inflammation. Post mortem biopsies in patients that died from COVID-19 point to airways and lungs as primary targets of the disease (11, 12) with advanced diffuse alveolar damage, pulmonary thrombosis and abnormal syncytia formation (13, 14). Several studies suggest that cytokine storm and inflammatory infiltrates in the alveolar space are associated with disease severity and death in COVID-19 (15, 16). While SARS-CoV-2 is genetically close to SARS-CoV, it shows much higher effective transmissibility (17, 18). One reason for this higher contagiousness is an active virus replication in tissues of the upper respiratory tract at an early stage of infection, with a high number of virus copies produced four days after the beginning of symptoms, and an active replication in the throat (19) (20). Furthermore, Zou et al (21) reported that the viral load detected in asymptomatic patients was similar to that of symptomatic patients on day 4 after symptoms onset, suggesting equal transmission potential of asymptomatic or minimally symptomatic patients at very early stages of infection (22). Epidemiological data have demonstrated that if all ages of the population are susceptible to SARS-CoV-2 infection, SARS-CoV-2 infection severity is different between children *versus* the adult populations and varies with age (23). A recent multi-national epidemiologic study found that children under 9 years old have very low case-fatality rates of SARS-CoV-2 infection compared to older patients (24). Moreover, these studies consolidate a large discrepancy in death rates of SARS-CoV-2 infected patients associated with age. Death rate in children (<9 years) is under 0.001% increasing to 8% in elderly patients (>80 years). A recent metadata analysis of several studies came to the same correlation between age and severity (25). The reason for this age-related discrepancy is not clear and could be linked to a decreased transmission and/or viral load with SARS-CoV-2 in children compared to adults. Only limited data are available about the mechanism of viral spreading over time and how the virus is released from the epithelia and might participate in the transmission of the infection between individuals or within an individual. Over the course of a 51 days period, infection of a reconstituted human airway epithelium infected with SARS-CoV-2 showed multiple waves of viral replication associated with a degradation of tight junction and a decrease in ciliary expression (26, 27). In this model, plaque-like cytopathic effects could be observed with the formation of multi-nucleated cells (28). Regarding the inflammatory response, interferon induction appears limited in the most severe clinical cases (29–31). In contrast, release of INF-λ was induced at day 4 post-infection of bronchial epithelia (BE). Noteworthy, viral RNA production in BE increased at day 2, suggesting a delay in the induction of the cellular antiviral response. A very recent report studying cell-intrinsic changes occurring in differentiated human nasal epithelial cultures from children, adults and elderly, have shown that ageing contributed to viral load, transcriptional responses, IFN signaling and antiviral responses (32). Yet, such data using a model mimicking the human bronchial epithelium are still missing. Here, we developed a model of reconstituted bronchial epithelium (BE) in air-liquid interface derived from bronchial epithelium samples of adult donors, which is the primary site of SARS-CoV-2 infection. We monitored the replication of SARS-CoV-2 over several days and followed virus spread in the epithelia. Using high-resolution imaging, we observed the massive formation and apical release of syncytia occurring between day three and four post-infection. We showed that syncytia and cells released into the apical lumen are infectious, suggesting they contribute to the spreading of the virus in the epithelium, and by extension, may transmit virus within the patient to the lower respiratory tract or into the environment. Furthermore, using reconstituted BE derived from children, we showed that viral production in children epithelia is very low compared to adults, and that viral spread is restricted. These results may explain the clinical and epidemiological observations that SARS-CoV-2 is more likely to infect older patients than children and that older patients show more severe clinical manifestations.

## Results

### Generation of a fully differentiated bronchial epithelia model

One of the major initial targets for SARS-CoV-2 is the respiratory tract. Primary infections often initiate in the upper respiratory tract from which they can spread to the lower respiratory tract to cause severe disease (18). Bronchial epithelia are pseudo-stratified cell layers with typical cell junctions, as well as a mucus layer and beating cilia on the lumen side (33, 34). To study the SARS-CoV-2 infection process in a physiologically relevant model, we established a cellular *in vitro* model of bronchial epithelia differentiated in air-liquid interface from individual donors (Fig. S1). Primary bronchial epithelial cells were collected from surgical bronchial resection or fibroscopy from individual adult donors at the Bordeaux university hospital. Patients were between 46 and 63 years old with a normal body mass index [BMI] (Table 1). Basal epithelial cells were expanded *in vitro* in culture flask until confluence. Basal cells were then seeded on cell culture insert and differentiated at the air-liquid interface for approximately 21 days (Fig. S1A). Using this differentiation protocol, we were able to generate between 12-24 individual inserts from a single donor allowing comparative analysis. Immuno-fluorescence (IF) analysis confirmed the presence of differentiated cell types. Specific antibodies allowed the detection of acetylated tubulin and mucin, characteristic of multi-ciliated cells and goblet cells respectively (Fig. S1B, movie S1) or acetylated tubulin and cytokeratin 5 (multi-ciliated cells and basal cells, Fig. S1C, movie S2). This analysis confirmed the pseudostratified apical-to-basolateral organizational integrity of the epithelia, *e.g*. a single cell layer of apical multi-ciliated cells covering a layer of basal cells and was further confirmed by electron microscopy (Fig. S1D). The presence of well differentiated cilia structures and tight junctions was also confirmed (Fig. S1D). Next, we determined the localization of ACE2, the primary receptor for SARS-CoV-2 in our model using IF analysis (Fig. S1E, movie S3). Co-label with antibodies against ACE2 and acetylated tubulin confirmed that ACE2 was expressed in apical multi-ciliated cells as previously reported (4, 35). Moreover, our data showed a prominent exposure of ACE2 on individual cilia reaching into the apical lumen (orange arrows), which suggests facilitated access *e.g*. for virus coming in through the respiratory tract.

**Table 1:** Clinical characteristics of adults and children donors used in this study. F: Female: M: Male; BMI: Body Mass Index.

| Patients | Sexe | Age | BMI |
| --- | --- | --- | --- |
| Adult A1 | F | 46 | 23 |
| Adult A2 | M | 58 | 25 |
| Adult A3 | F | 54 | 24 |
| Adult A4 | M | 63 | 25 |
| Adult A5 | M | 51 | 25 |
| Adult A6 | F | 61 | 21 |
| Child C1 | F | 13 | 17 |
| Child C2 | M | 12 | 16 |
| Child C3 | F | 12 | 18 |

### SARS-CoV-2 monitoring and BE infection

Next, BE were inoculated on the apical side with a suspension of a reference SARS-CoV-2 strain (BetaCoV/France/IDF0372/2020) at a multiplicity of infection (MOI) of 0.012. Apical and basolateral compartments were collected 3 days post-infection (dpi) and used to infect Vero E6 cells (Fig. 1A). A cytopathic effect (CPE) was observed in the Vero E6 cell culture as early as 2 days post-infection when inoculated with the apical washes, indicating an effective infection and replication of the virus (Fig. 1A). When using the basal medium, 3 days of inoculation were necessary to observe a similar CPE (Fig. 1A). This faster appearance of CPE when using the apical fraction may be correlated to a higher viral titre compared to the basal medium. To ascertain that this CPE is indeed due to viral replication and not a toxic effect from the inoculation, we extracted total RNAs from the Vero E6 supernatant on the next day (4 dpi) and quantified viral RNAs using in-house qRT-PCR targeting the N-gene region. No RNA could be detected in the supernatant of Vero E6 cells inoculated with either the apical or basolateral fractions obtained from non-infected BE (Fig. 1B, control). In contrast, when using basolateral or the apical fraction from infected BEs, the Vero E6 supernatant contained high level of SARS-CoV-2 RNA, comparable to what is observed with a direct infection of Vero E6 cells infected at a MOI of 0.01 (Fig. 1B). These data attest that SARS-CoV-2 actively replicates in reconstituted BE and that inoculation from the apical side results in an active infection. To detect virus-infected cells, we generated monoclonal antibodies against the SARS-CoV-2 N nucleocapsid protein using bacterially expressed and purified full-length protein as detailed in the methods section. Hybridoma supernatants were tested using western blot and IF detection through confocal microscopy (Fig. S2). Of several positive clones, hybridoma clone 3G9 was selected for this study as it specifically recognized the N protein of SARS-CoV-2 (Fig. S2A) and detected infected cells in IF staining (Fig. S2B). To investigate which cell type is the primary target during SARS-CoV-2 infection, fully differentiated epithelia were infected with SARS-CoV-2 at a MOI of 0.01 for 1 h from the apical side after which the viral suspension was removed. Epithelia were fixed 24h post-infection in 4% paraformaldehyde (PFA) and processed for IF analysis using SARS-CoV-2-N specific antibodies. We successfully detected infected cells in the BE (green signal Fig. 1C-E). Specific co-label of Muc5A showed that goblet cells were not infected (magenta signal, Fig. 1C, movie S4). Similarly, no co-localization could be observed between the SARS-CoV-2 N protein and CytK5 showing that basal cells were not infected either (Fig. 1D, movie S5). Conversely, the signal arising for the N protein staining was systematically associated with strong labelling for acetylated tubulin, a specific marker for multi-ciliated cells (orange arrow, Fig. 1E, movie S6). This is consistent with previous reports that apical multi-ciliated cells are the primary target cells for SARS-CoV-2 infection (4, 28, 36). In addition, all BEs were co-labelled with fluorescent phalloidin to mark cell boundaries for 3D imaging of the entire epithelial depth. Infected cells were exclusively located at the apical surface of the BE (Fig. 1C-E). All IF data were confirmed using BE generated from at least two different donors, suggesting that the primary infection of epithelial cells is determined by the epithelia architecture and is not due to the genetic background of the donor.

**Figure 1.**
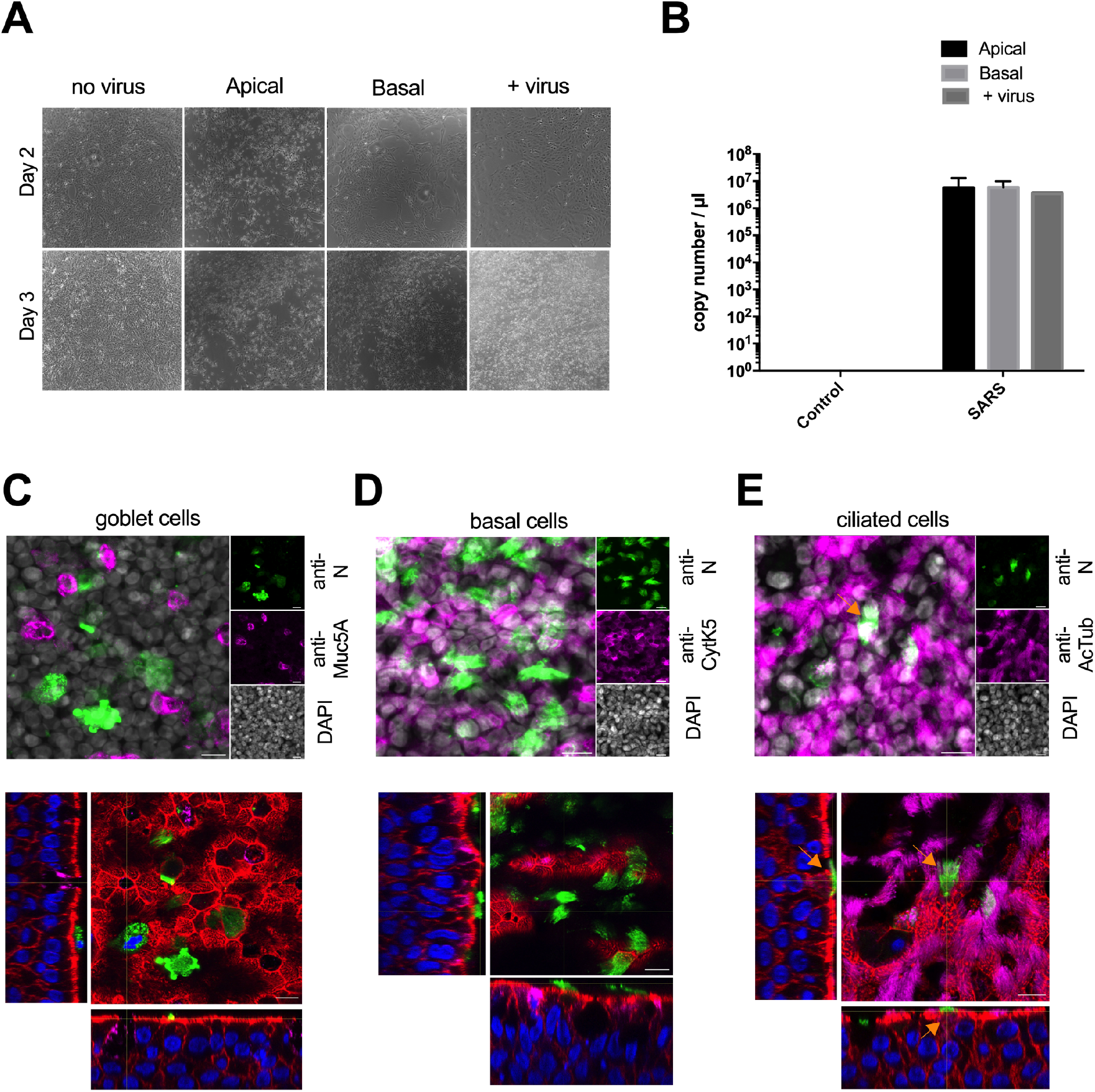
SARS-CoV-2 infection of bronchial epithelia (BE). A: The left panel shows brightfield microscopy images of Vero E6 cells at either day 2 (top row) or day 3 (bottom row) post-inoculation with apical washes, basolateral media or a viral suspension (MOI of 0.01). B: Quantification of SARS-CoV-2 RNA in Vero E6 supernatant. Total RNA were extracted 96h post-infection and quantified by qRT-PCR. Mean and standard deviation are derived from 3 independent determinations except for the positive control (Vero E6 infected directly with a viral suspension instead of BE fractions). C: Differentiated BE were infected with SARS-CoV-2 and stained 24h post-infection with anti-N (green signal) to identify infected cells and anti-Muc5A to detect goblet cells (magenta signal) and counterstained with DAPI (grey signal). Top image shows a Z-projection, the bottom image shows an individual Z-section of a 3D reconstruction counterstained with phalloidin. Scale bar is 10μm, for full Z-stack see movie S4. D: Experiment and presentation as in (B) stained with anti-N (green signal) to identify infected cells and anti-cytokeratin 5 to identify basal cells (magenta signal) and counterstained with DAPI (grey signal). Scale bar is 10μm, for full Z-stack see movie S5. E: Experiment and presentation as in (B) stained with anti-N (green signal) to identify infected cells and anti-acetylated tubulin to identify multiciliated cells (magenta signal) and counterstained with DAPI (grey signal). Scale bar is 10μm, for full Z-stack see movie S6.

### Infection kinetic of epithelia from different adult donors

To better understand how SARS-CoV-2 spreads in the epithelium after initial infection of multi-ciliated cells, we infected BEs from four individual adult donors (A1 to A4, Table 1) and monitored them over the course of 7 days. Low magnification images obtained using IF microscopy showed that N protein could be detected within 24h of infection in a small number of cells (Fig. 2A). Nonetheless, the signal number and intensity increased drastically from the 2 dpi time-point and tended to decrease slightly towards the end of the observation period (Fig. 2A). Similar results were obtained with the other two donors suggesting rapid onset of viral replication and spread (not shown). We quantified the number of N-positive signals at low resolution for each donor confirming that the number of infected cells strongly increased within two to three days of the initial infection, reaching a maximum around day four, and consistently decreased somewhat on the seventh day for all donors (Fig. 2B). Of note, much larger N protein associated signals could be observed at the peak of the infection. These larger structures were co-labelled with cytokeratin 5, the marker for basal cells (see arrows in Fig. 2A). This observation started on the third day but was most prominent on the fourth day and was observed for all donors. Therefore, we also quantified the size of the N protein associated signals over time (Fig. 2C). The analysis revealed a statistically significant average increase in signal size between the third and fourth day for all four donors. In parallel to the imaging analysis, release of newly produced viruses into the apical mucus was quantified by qRT-PCR (Fig. 2D). For all four donors, the viral RNA copy number correlated with the observed cellular N protein labelling with a fast increase from day 2 reaching a plateau between 3 and 4 dpi. Altogether, these dat a suggested that apical SARS-CoV-2 inoculation of BEs resulted in efficient infection and subsequent progeny production and release into the apical lumen.

**Figure 2.**
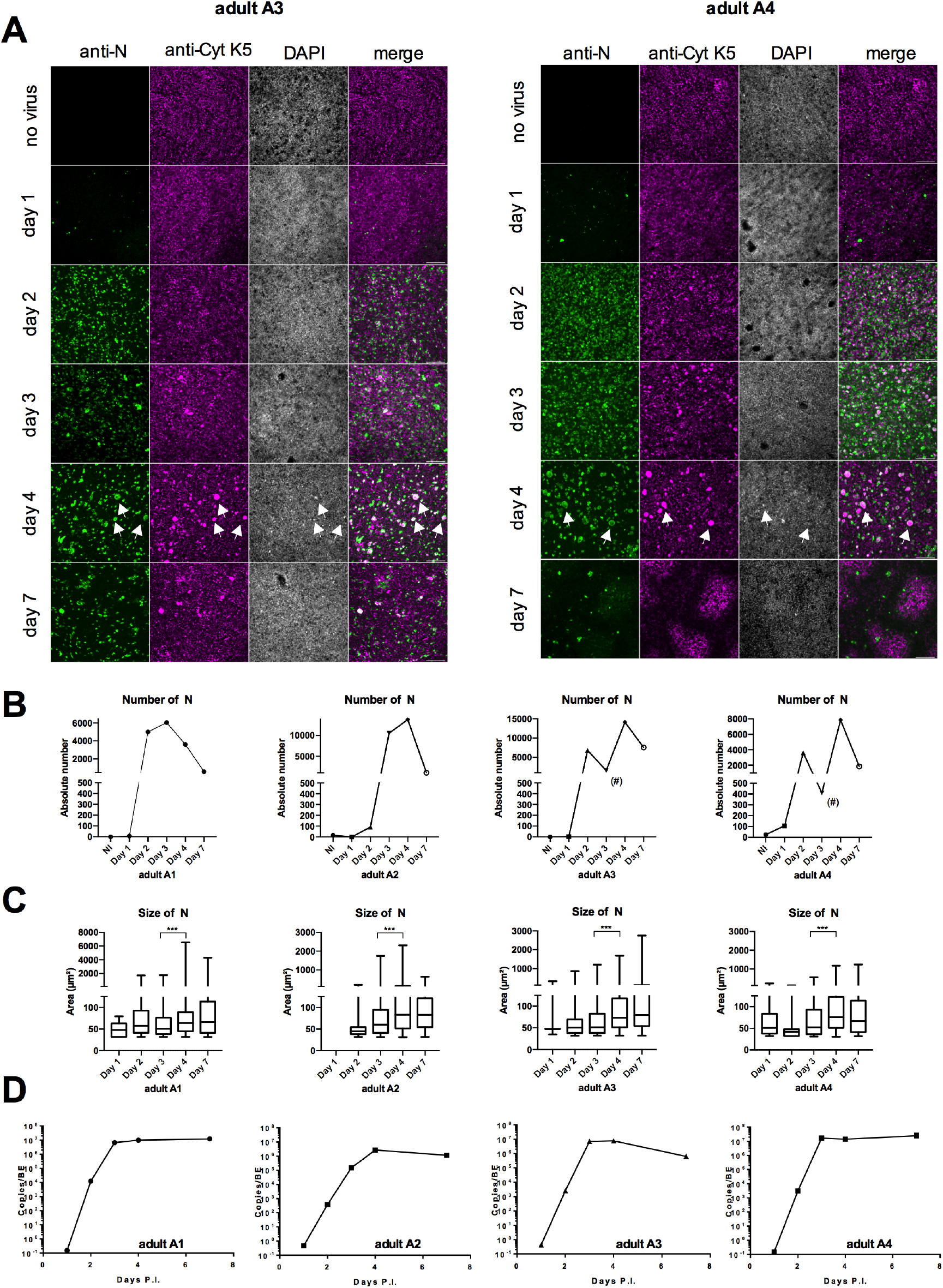
SARS-CoV-2 infection kinetic of bronchial epithelia (BE). A: Representative widefield microscopy images of BE from two adult donors (A3 left panel, A4 right panel) at low resolution. Scale bar is 20μm. BE were fixed at day 1, 2, 3, 4, 7 as indicated to the left of each row, non-infected controls were also fixed at day 7. BE were stained with anti-N antibodies to detect infected cells (green signal first column), anti-cytokeratin 5 to detect basal cells (magenta signal, second column) and counterstained with DAPI (grey signal, third column) and a merge of the three signals (forth column). Large specific signals in all channels are apparent on day four (white arrows). B: The absolute number of N-positive signals was determined for each BE for the whole epithelia on each day as indicated. Data shown are absolute number of N dots quantification at different days post-infection and described in material and methods. The (#) sign marks points with partial BE damage C: Signals quantified in (B) were classed by size and plotted as min to max Box & Whisker plots, ***: P < 0.001 based on One way ANOVA. D: Apical washes for each BE were subject to RT-qPCR analysis to determine genome copy numbers at day 1, 2, 3, 4, and 7 post-infection as indicated.

### Infected multi-ciliated cells form syncytia with basal cells at the apical side of the BE

Using high-resolution microscopy, we observed that larger N-positive signals corresponded to multinucleated cellular structures reminiscent of syncytia. These syncytia could be found in all regions of the epithelia (Fig. 3) and their formation at day 4 was common to all four donors tested. In contrast, we did not observe any syncytia formation in non-infected control epithelia. Unexpectedly, the N-positive syncytia forming on day three and four also stained positive for the basal cell marker cytokeratin 5 (Fig. 3A). This was not the case at earlier time points (day one and two) where basal cells rarely stained positive for N protein and did not form syncytia. Accordingly, we quantified the number of double positive syncytia (*i.e*, nucleocapsid protein and cytokeratin 5) over time (Fig. 3B). The proportion of double positive cells (*i.e*, syncytia) increased constantly and reached a maximum on the fourth day after which there is a drastic drop in double positive cells (Fig. 3B, upper panel). Normalization of the double positive cells for either the total amount of basal cells (Fig. 3B, middle panel) or the total amount of infected cells (Fig. 3B, lower panel) revealed that double positive cells but not overall infected cells disappeared on day four. We also observed that the newly formed multinucleated cells only partially stained for acetylated tubulin (Fig. 3C). Zooming in on different regions of the epithelia revealed that newly formed syncytia frequently lost their stain for acetylated tubulin (Fig. 3C side panel). Moreover, syncytia that still expressed acetylated tubulin presented an amorphous staining, and rarely distinguished cilia features. Similarly, part of the syncytial structures failed to stain with phalloidin (*e.g*. Fig. 3A, right panel), that was used to delineate cells in the epithelia. Altogether, Cytokeratin 5, phalloidin and acetylated tubulin staining patterns suggested that syncytia were formed through the fusion of infected ciliated cells with basal cells, associated with the loss of cilia and reorganization of cytoskeletal features including the actin and tubulin cytoskeleton. Furthermore, three-dimensional imaging of epithelia showed that syncytia formed exclusively on the apical side of the epithelium and forming an elevated layer on top of the epithelia (Fig. 3A right and 3C bottom panel, see also movie S7 and S8). These extrusions were also observed using EM (Fig. 3D, white asterisk), but never in the context of non-infected epithelia (Fig. S1). These structures harbored only reminiscent cilia structures in place of multi-ciliated cells in non-infected epithelia. This latter observation is consistent with previous reports showing that SARS-CoV-2 infection of lung epithelial cells trigger the partial loss of cilia (27). Importantly, using EM we observe vesicular inclusions within those extruded cells that contained virus particles indicating that multi-nucleated infected structures actively produced viruses (Fig. 3D, black arrows).

**Figure 3.**
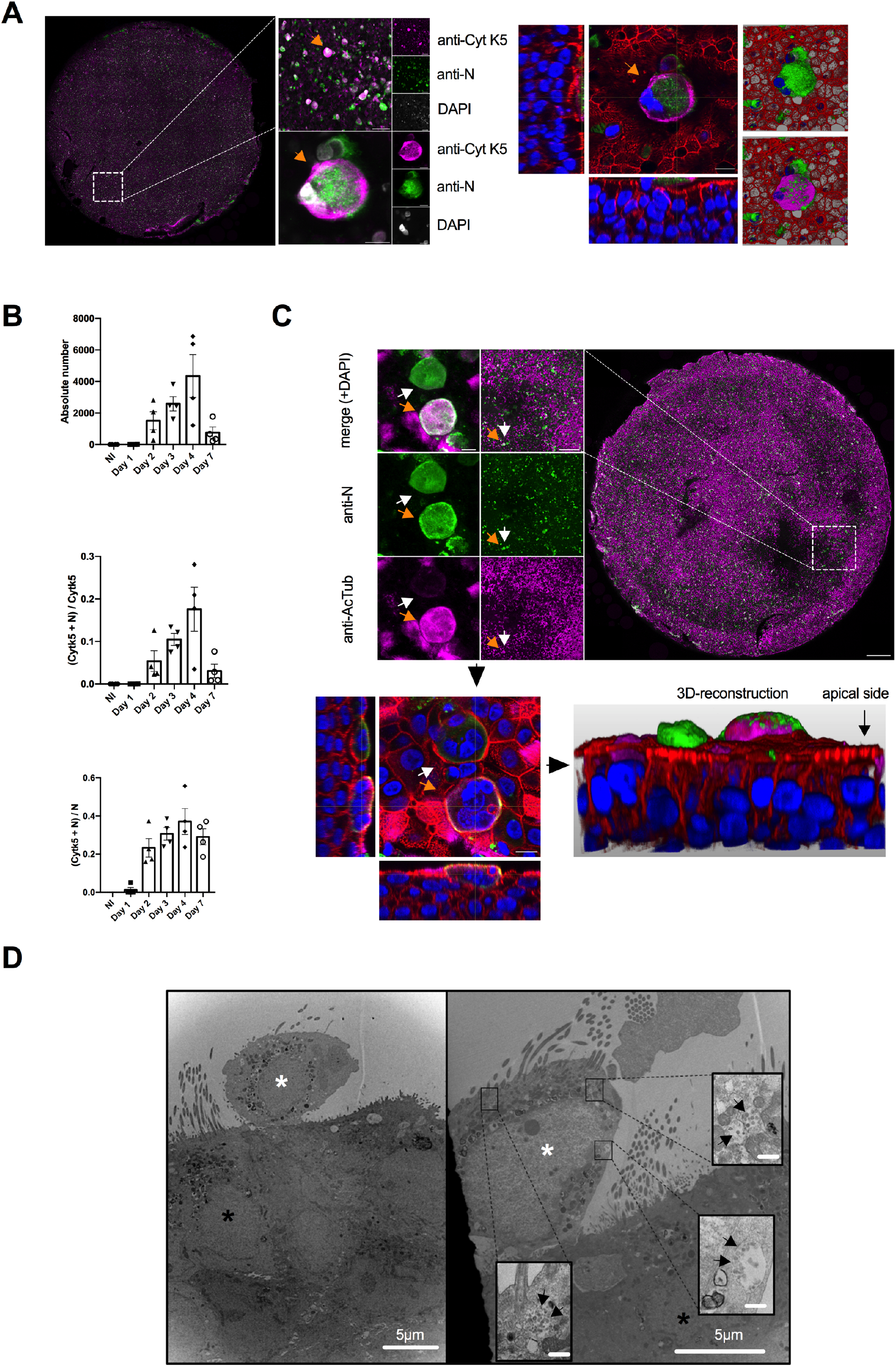
SARS-CoV-2 infection of bronchial epithelia (BE) induces apical syncytia. A: High resolution image analysis of entire BE 4 days post-infection (left panel, adult A4). BE were stained with anti-N antibodies to detect infected cells (green signal) and anti-cytokeratin 5 marking basal cells (magenta signal) and counterstained with DAPI (grey signal in the left panel, blue on the right panel). The boxed inset is magnified and one individual syncytia is indicated by the orange arrow. The syncytia is further magnified as maximum Z-projection (left) or as individual Z-stack (right, with phalloidin counterstain in red), or as 3D image reconstruction to see its apical location. Scale bar is 10μm, 50μm, 200μm respectively. See also movie S7. B: Estimation of the total number of double positive (N and cytokeratin 5) was determined (top panel) and normalized for total number of basal cells (middle panel) and total number of infected cells (bottom panel). Data shown are absolute number of colocalisations between N and cytokeratin 5 for four different donors, as determined using semi-automatic quantification. Absolute number of colocalisation was then normalized by absolute number of cytokeratin 5 or N positive cells. Data are presented as mean ± SD, n = 4. C: High resolution image analysis of entire BE 4 days post-infection. Experiment and image representation as in (A). BE were stained with anti-N antibodies to detect infected cells (green signal) and anti-acetylated tubulin detecting multi-ciliated cells (magenta signal) and counterstained with DAPI (grey signal in the top panel, blue on the bottom panel). Double positive syncytia are marked by orange arrow, single positive syncytia with white arrow. Scale bar is 10μm, 50μm, 200μm respectively. See also movie S8. D: Electron micrograph of infected BE 4 days post-infection. The large images show extruded cell on the apical side of the epithelia (white asterisk) adjacent to multi ciliated cells (black asterisk). The insets show virus containing vacuoles in the extruded cell as indicated by black arrows. Scale bars are provided in the image (5μm for large images and 200nm for insert images).

### Infected cells and syncytia are released into the apical BE lumen and transmit infection

Because cells and syncytia were extruding from the epithelium, we wondered whether infected cells/syncytia could be released from the epithelium and account for the spreading of the infection. To test this hypothesis, we infected epithelia from two donors for three and four days. Apical washes of epithelia were performed after three and four days of infection and concentrated on microscope slides *via* cytospin. After IF processing, we showed that apical washes contained individual infected cells (positive for N-protein staining) but also several infected syncytia, suggesting that both are indeed released into the apical epithelial lumen (Fig. 4A). To test the relative infectivity, apical washes were clarified of cell material by low-speed centrifugation. Both the clarified supernatant and the removed cellular fraction were used to infect Vero E6 cells. After 24h, cells were fixed and analyzed by IF microscopy. Inoculation with the supernatant as well as the cellular fraction of the apical wash showed efficient Vero E6 cell infection (data not shown). In parallel, inoculated Vero E6 cells were monitored for the appearance of a virus-induced CPE. Apical wash after 4 days of epithelia infection resulted in CPE within 48h whereas a comparable CPE required 72h with an apical wash resulting from a 3 days infection (data not shown). AT 96h post-infection the Vero E6 cell CPE was quantified (Fig. 4B).

**Figure 4.**
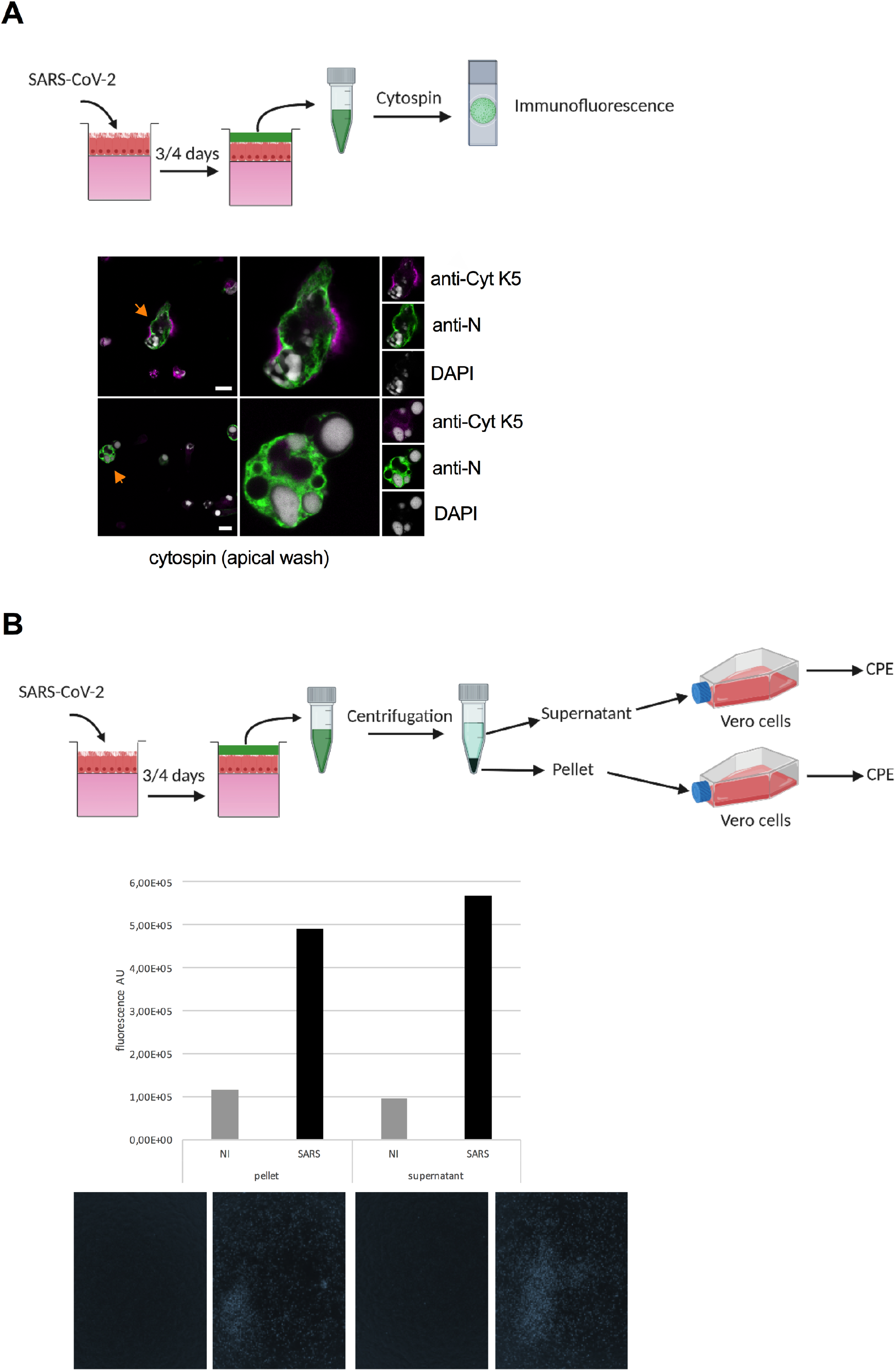
The apical bronchial epithelia lumen contains infected cells that transmit infection. A: A schematic of the experimental design (top panel). Apical washes at 4 days post-infection were fixed and concentrated on slides using cytospin (bottom panel) and stained with anti-N antibodies to detect infected cells (green signal) and anti-cytokeratin 5 marking basal cells (magenta signal) and counterstained with DAPI (grey signal). Arrows indicate syncytia in the overview and are magnified to the left. The boxed inset is magnified and individual syncytia are indicated by the orange arrows. Scale bar is 10μm. B. A schematic of the experimental design (top panel). Apical washes were separated into supernatant and cell pellet and used to infect Vero E6 cells. CPE was quantified at 96h post-infection using fluorescence readout as described in material and methods (Bottom panel).

Apical washes from non-infected epithelia produced only background levels of cell death (gray bars, Fig. 4B). In contrast, significant levels of cell death occurred in Vero E6 cell after inoculation with either the supernatant or the pellet (cellular fraction) of an apical wash issued from an infected epithelium (black bars, Fig. 4B). Taken together this analysis shows that epithelia produce and release large amounts of new viruses into the apical lumen, a significant fraction of the released infectious virus dose stems from infected cells and syncytia.

### Epithelia from children partially restrict SARS-CoV-2 infection but not syncytia formation

In our experimental model, each epithelium can be traced to an individual donor, while generating enough individual inserts to allow biological repeats and kinetic studies. Our analysis showed that epithelia from several adult donors responded similarly to the infection with SARS-CoV-2. A striking observation was that infections spread very fast over the entire epithelia and produced vast amounts of syncytia for apical release in a synchronized manner. Adult donors in this study were between 46 and 63 years old (table 1), which puts them statistically into a medium/high risk group to develop severe COVID-19 symptoms. In contrast, several reports have indicated that children are much less susceptible to severe forms of COVID-19, while their role in spreading virus infections is controversially discussed (37, 38). To investigate whether SARS-CoV-2 infects BEs differently depending on the age of donors, we prepared epithelia through expansion and differentiation of bronchial epithelial cells obtained from children (Table 1) that have undergone bronchial fibroscopy for chronic bronchopathy (child C1) or bronchiectasis (children C2 and C3). Fully differentiated epithelia from children showed the same cellular arrangement (epithelial cells, basal cells, goblet cells) and physiological properties (cilia beating, mucus production) as adult derived epithelia. A kinetic experiment was performed to compare the SARS-CoV-2 infection dynamics in BEs derived from children (C1 to C3) or from adult donors (A5 and A6). The BEs were fixed at 1, 2, 3, 4 and 7 d.p.i. with a non-infected control for each donor run in parallel and fixed at day 7. Individual epithelia were fixed and processed for IF analysis using antibodies against cytokeratin 5, SARS-CoV-2 N-protein and counterstained with fluorescently labelled phalloidin and DAPI. As observed before (Fig. 2), infecting BEs from adult donors at a MOI of 0.012 resulted in a fast increase in the presence of infected cells (within 48h) and the formation of a significant amount of syncytia on the fourth day (A6, Fig. S3A). In sharp contrast, all child derived epithelia showed a remarkable resistance to virus infection (Fig. S3 B-D). Of note, virus spread differed significantly in BE originating from the individual child donor. A slow but substantial increase in infected cells over time was observed in BE derived from donor C1 (Fig. S3B). In comparison, BE derived from C2 did not support substantially increase of the number of infected cells after the initial appearance of positive cells (Fig. S3C) and BE derived from the last donor, C3, only ever showed very few infected cells, reminiscent of an abortive infection (Fig. S3D). Low magnification imaging of the entire epithelia showed that initial infections in BE from donor C1 were limited to few cells. The N-protein associated signal seemed to grow over time into foci of infection that further enlarged by infecting surrounding cells at the periphery (Fig. 5A, top row, left). High-resolution images confirmed that cells at the foci border stained strongly, while several cells surrounding these foci were already positive for SARS-CoV-2 N-protein. This suggested the existence of a front of highly replicating cells with forward cell-to-cell or short range spread as infection mode (Fig. 5B). In contrast, infection spread in epithelia from donor C2 seemed to be even more restricted (Fig. 5A, bottom row, left). In the case of the BE derived from the child donor C2, most of the cells that were initially infected at day one/two developed into local cluster of infected cells without much lateral spread. High-resolution imaging revealed that within these clusters, several cells fused with basal cells to form small syncytia that had apical localization (Fig. 5C, movie S9), reminiscent with what was observed in adult donors. It is only after seven days of infection that some spreading into small patches could be observed, mimicking observations made for C1 on the second and third day of infection. Low magnification imaging of the entire epithelia derived from child C3 confirmed sporadic infection signals in the BEs, while the adult-derived control A7 showed massive spread of the infection throughout the entire epithelia at 4 d.p.i (right panel, Fig. 5A). Quantifying the total number of infected cells in each epithelium confirmed our observation (Fig. 5D). In contrast to the adult control, no significant differences in signal size was observed between day three and day four for either of the children derived epithelia (Fig. 5E). Still, the average signal size appeared larger likely due to clustering of infected cells (Fig. 5E). For all three children derived epithelia and the adult control we also measured the accumulation of SARS-CoV-2 in the apical lumen using quantitative PCR (Fig. 5F). The quantities of released virus over time accurately reflected the spread of infection observed by IF and quantification of infected cells. Taken together our analysis clearly demonstrated that epithelia from children were less susceptible to SARS-CoV-2 and exhibited an intrinsic resistance towards virus infection and/or spread. One possible explanation for this intrinsic resistance of children BE could be differences in IFN response (39, 40) or morphological differences (41). Accordingly, we compared the accumulation of interferon λ 1/3 and measured the concentration in BE medium from adults and children in response to SARS-CoV-2 infection (Fig. 5G). We did not find interferon λ at the beginning of the infection. Children BE secreted interferon λ starting at day 1 post-infection whereas adult BE produced detectable amount of interferon λ only at day 3 post-infection. Interferon λ concentration increased subsequently for both age groups and reached similar levels at day 4 and 7 post-infection (Fig. 5G). The difference in the kinetic for Interferon l secretion between adults and children in response to SARS-CoV-2 infection may thus provide an explanation why in our model children derived BE resist better to SARS-CoV-2 infection. Still, the strength of this resistance differs from donor to donor and delays virus spread to different degrees or may prevent virus spread entirely.

**Figure 5.**
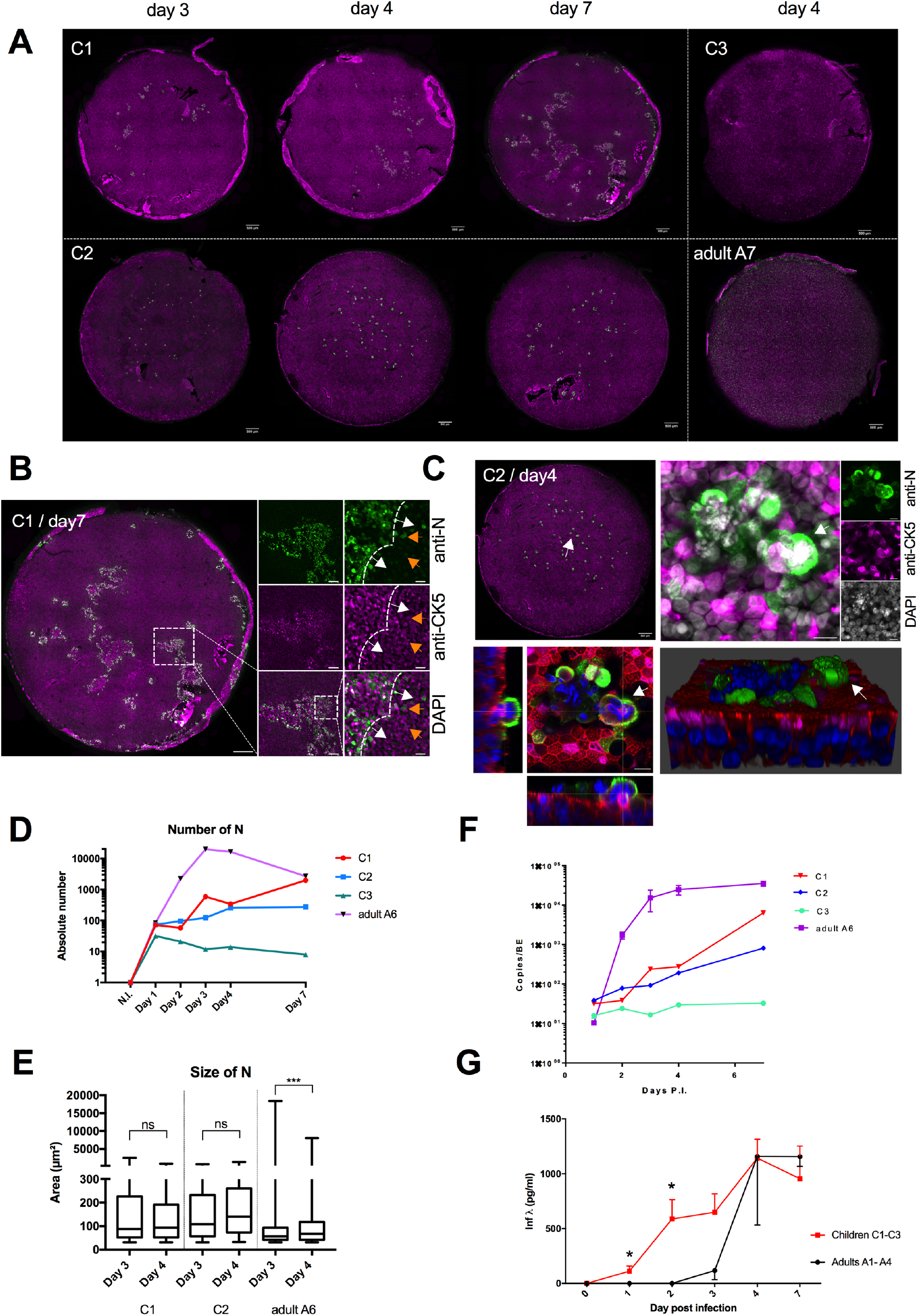
SARS-CoV-2 infection of bronchial epithelia (BE) in children. A: Overview of entire BE from children and adult control. Donor are as listed in table 1 (C = child, A = adult control) and days post-infection is indicated on the top. BE were stained with anti-N antibodies to detect infected cells (green signal) and anti-cytokeratin 5 marking basal cells (magenta signal). B: High resolution imaging of the Child C1 epithelia 7 days post-infection. Epithelia were stained with anti-N antibodies to detect infected cells (green signal) and anti-cytokeratin 5 marking basal cells (magenta signal). The boxed area containing an infection foci is magnified to the right. The higher magnification shows the infection front (dashed line and white arrows) and individual infected cells in the vicinity of the infection front (orange arrows). C: High resolution imaging of the Child C2 epithelia 4 days post-infection. Epithelia were stained with anti-N antibodies to detect infected cells (green signal) and anti-cytokeratin 5 marking basal cells (magenta signal) and counterstained with DAPi (grey or blue signal). The white arrow in the overview points at a syncytium that is further magnified as maximum Z-projection (left panel) or individual Z-stack (right panel with phalloidin counterstain in red), or as 3D image reconstruction to see its apical location. Scale bar is 10μm, 50μm, 200μm respectively. See also movie S9. D: The absolute number of N-positive cells throughout the BE was estimated for three children donor (C1-3) and one adult control (A6). Color code as indicated. E: The average size of N-positive signals from (D) was determined for two child epithelia (C1 and C2) and one adult control (A6) at 3 and 4 days post-infection as indicated. F: Apical washes for each BE were subject to RT-qPCR analysis to determine genome copy numbers at day 1, 2, 3, 4, and 7 post-infection as indicated. Color code same as in (D). G: Basolateral supernatant from adults (black, n=4) and children (red, n=3) BE were subject to ELISA to determine interferon λ concentration at day 0, 1, 2, 3, 4 and 7 post-infection as indicated. Results are presented as mean ± SEM and * indicates significant difference between adults and children for a time point using Mann Whitney t test.

## Discussion

In this study, we used human reconstituted bronchial epithelia to investigate the onset of infection and replication of SARS-CoV-2 in BE. Our approach uses primary cells from individual patients obtained through our local hospital collected in the bronchial tree between the third and fifth generation. Bronchial epithelia are an important tissue to study because following initial infection in the upper airways, subsequent infection of the bronchial tissue determines whether a SARS-CoV-2 infection results in severe or mild respiratory illness by controlling the spread into the lower respiratory tract. These features make a distinction in our approach from similar studies using primary respiratory cells from either upper airway (nasal, tracheal) (3, 26) or commercial sources (27) with undefined donor material. Using this physiological model, we observed SARS-CoV-2 production mainly on the apical side of the epithelia following infection. We then used immunofluorescence imaging to follow and compare the infection in the BE from several individual donors for seven days. This approach allowed the detection of infected cells as early as 24h post-infection. The infection spread throughout the whole epithelia within three to four days post inoculation followed by a drop in the number of infected cells on the last day. Quantification of viral RNA confirmed these observations and showed that viral replication reached a plateau around day four post-infection. Using cell specific markers, we identified that infected cells during the first two days corresponded largely to multi-ciliated cells staining positive for acetylated tubulin in agreement with previous studies (26, 28). We rarely observed infected basal or goblet cells during this period. Interestingly, starting at day three of the infection not only the infected cell number but also the signal size of infected cells increased with a statistically significant shift towards larger cells between day three and four. We could show that the larger signals corresponded to infected multinucleated syncytia. The formation of cell fusions in coronavirus infected primary airway epithelia was previously reported (28, 42) but not systematically detected (26, 27, 32). In our study, we were able to find extensive syncytia formation in all six adult donors. Syncytia formation was transient and reached a maximum at day four to sharply drop towards the seventh day. The fusogenic potential of SARS-CoV-2 is well known and involves the Spike protein and the ACE2 receptor (43, 44). When we used cell specific markers to identify the syncytia cell composition we found that several syncytia were double positive for the basal cell marker cytokeratin 5 as well as the cilia marker acetylated tubulin or only for the basal cell marker (Fig. 3). This suggested that syncytia are formed by the fusion of basal cells with infected multi-ciliated cells. This is consistent with previous reports that infection of multi-ciliated cells with SARS-CoV-2 results in cilia loss and cell dedifferentiation (27, 28). Fusion of initially infected multi-ciliated cells with basal cells as one mode of virus cell-to-cell spread was further supported by quantification of infected syncytia positive for cytokeratin 5, which constantly increased in number until day four in all analyzed donors. The sharp drop on day four in the number of double positive syncytia, but not in the number of overall infected cells, is consistent with our observation that syncytia were extruded at the apical side of the epithelia. We found frequent syncytia forming at the apical side of the epithelia positive for the N nucleoprotein and EM analysis showed that they indeed contained high amounts of virus trapped in a vesicular compartment. Furthermore, we were able to show that infected syncytia are released into the apical supernatant and that released syncytia and infected cells are as infectious as free virus (Fig. 4). This strongly suggests that infected syncytia and cell release into the apical lumen could be an important contribution to the spreading of large and compact amounts of viruses into the upper respiratory tract from which cell associated virus can either decent into the lower respiratory tract or reach the environment increasing the actual infectious dose. Interestingly, pathology reports from patients succumbed to Covid-19 show abnormal syncytia formed by pneumocytes in the lower respiratory tract suggesting that our observations in the BE model find their counterpart in severe forms of COVID-19 (14, 45, 46). Accordingly, such an event would be in agreement with the clinical observation of hospitalized patients, which reported a high detection of SARS-CoV-2 in sputum and its transmission by droplets (19). The fact that syncytia production is massive but transient is well correlated with another report showing that virus production in a primary airway epithelium is cyclic with peaks of virus release every 7-10 days (26). The authors suggest that this periodicity is driven by recurrent epithelia removal and regeneration. Interestingly, such a peak in virus production would provide an explanation for the phenomenon of “super spreader”, frequently suggested based on epidemiological data (47). A periodicity or variability in the quantity of virus released from infected tissue thus may affect contagion. Furthermore, we also observed the loss of cilia in many of the syncytia, which could be responsible for a poor mucociliary clearance that impedes the evacuation of viral particles and pathogens. Taken together, our findings are in accordance with previous findings but highlight syncytium formation as an important mechanism to explain the spreading of SARS-CoV-2 and the physiopathology of bronchial epithelium infection (14, 43, 45, 46). Since the beginning of the COVID-19 pandemic, SARS-CoV-2 infection is more virulent in adults compared to children. We explored the bronchial epithelium infection of children with SARS-CoV-2 and compared our observations with those made in BE from adult donors. Strikingly, we find very different spreading of SARS-CoV-2 in children BE *versus* adult BE. First, the overall viral production was very low in BE of children compared to adults, which is reflecting the slower kinetic in the onset of virus production over time. In agreement with the virus quantification, child epithelia showed a remarkable resistance to virus infection as very few infected cells were observed. Rather than rapidly spreading throughout the entire epithelia, as observed for adults, the infected cells in children form cluster or foci of infected cells. From these foci, the infection slowly spread into the surrounding bystander cells. Yet, syncytia formation was also observed, at least in one child, suggesting that the fusion of basal cells with multi-ciliated cells was not restricted to adult infected BE. However, the number of syncytia was much lower than in adults reflecting the low virus spread. The obvious difference in susceptibility to SARS-CoV-2 infection between adults and children, which we observed in the BE model is in agreement with the reduced epidemiological infection rate described for children and strong discrepancy in death rate between children and adults/elderly (24, 25). A very recent study using nasal BE also showed differences in the susceptibility to SARS-CoV-2 infection between adults and children (32).

The reason for this intrinsic difference between adults and children BE is unknown. One possible explanation could be an age-related variation in the expression or accessibility of the primary viral receptors (ACE2 and TMPRSS2) (48). We show that children BE have a quicker induction of interferon λ in response to SARS-CoV-2 infection starting as soon as 1-day post-infection whereas adults BE exhibit detectable level of interferon λ only 3-day post-infection. Recent studies show that SARS-CoV-2 blocks the interferon response by targeting the RIG-I/MDA-5 pathway (49, 50). Our study is consistent with a delay of Interferon λ production after SARS-CoV-2 infection in adults but less so in children. This could suggest that either SARS-CoV-2 is less efficient in counteracting the IFN response in children BE or alternatively, that the IFN response in children is faster and an antiviral state is induced throughout the epithelia that slows down virus spread. Such an age-related susceptibility of BE has been reported for other respiratory pathogens including respiratory viruses such as Rhinovirus-C, Adenovirus and RSV (Respiratory Syncytial Virus) (51, 52)(53) but also fungi (Aspergillus fumigatus) (54) and bacteria (Haemophilus influenzae) (55). Future studies will be required to study the exact mechanism behind the differences in IFN response that we observed. Taken together our data clearly demonstrate that BE from children are less susceptible to SARS-CoV-2 infection. Our data suggest that an accelerated interferon response might contribute to this resistance supporting timed interferon application as therapeutically beneficial concept in the treatment of SARS-CoV-2 infections (56–58).

## Materials and Methods

### Monoclonal antibodies and ethics statement

Monoclonal antibodies were raised against bacterially expressed and purified SARS-CoV-2 N protein in 3 mice using the protocol as previously described (59). Hybridomas were cloned by limiting dilution and screened by immunofluorescence on infected VERO cells. Clone 3G9 was retained for this study and antibody was affinity purified from hybridoma supernatant prior to use. Mice experiments have been performed in the conventional animal facilities of the University of Bordeaux (France) (approval number of B-33-036-917), with the approval of institutional guidelines determined by the local Ethical Committee of the University of Bordeaux and in conformity with the Ministry for Higher Education and Research and the French Committee of Genetic Engineering (approval number n °17621 -V5-2018112201234223).

### Viruses and cell lines

Vero E6 cells were maintained in Dulbecco’s modified Eagle’s medium (DMEM, Gibco) supplemented with 10% fetal calf serum (FCS) and gentamicin (50μg/mL) at 37°C in a humidified CO_2_ incubator. The SARS-CoV-2 strain BetaCoV/France/IDF0372/2020 was supplied by the National Reference Centre for Respiratory Viruses hosted by Pasteur Institute (Paris, France) through the European Virus Archive goes Global (EVAg platform). Agreement to work with infectious SARS-CoV-2 was obtained and all work with infectious SARS-CoV-2 was performed in a Class II Biosafety Cabinet under BSL-3 conditions at the UB’L3 facility (TBM core, Bordeaux).

### Viral production

The SARS-CoV-2 strain was produced by infecting Vero E6 cells at a multiplicity of infection (MOI) of 0.01, then incubating the cells at 37°C in a humidified CO_2_ incubator until appearance of a cytopathic effect (around 72 h). The culture supernatant was clarified by centrifugation (5 minutes at 1500 rpm) and aliquots were stored at −80°C. Stock titers were determined by adding serial dilutions to 2 × 10^4^ Vero E6 cells in supplemented DMEM in a 96-well plate. Eight replicates were performed. Plates were incubated at 37°C and examined for cytopathic effect. Quantification of cytopathic effect was determined using the Cell tox ^TM^ green cytotoxicity assay (Promega) according to manufacturer instructions and a Victor Nivo reader (Perkin Elmer). The TCID_50_ was calculated according to the method of Reed & Muench (60). PFU/ml was estimated from the TCID_50_ determination.

### Culture of primary bronchial epithelia (BE) and ethics statement

Bronchial epithelial cell culture was established from bronchial brushings or lung resection performed between the third and fifth bronchial generation from patients undergoing elective surgery as previously described (34). Bronchial epithelium explants were cultured using PneumaCult Ex medium (Stemcell, Vancouver, Canada) for expansion of basal epithelial cells at 37°C in 5% CO_2_. Then, 10^5^ basal cells were grown on cell culture inserts (Corning, New York, NY) within the air-liquid interface for 21 days using PneumaCult ALI medium (Stemcell, Vancouver, Canada). Such a culture allows the differentiation into pseudostratified muco-ciliary airway epithelium composed of ciliated cells, goblet cells, club cells and basal cells. The complete differentiation was assessed by the capacity of cilia to beat and mucus production under light microscope. The study received approval from the local and national ethics committee from the CNIL through the TUBE collections.

### Infection of epithelia

Prior to infection, epithelia were washed three times with PBS to remove mucus and basal ALI medium was exchanged with 500 μL of fresh medium. The inoculum containing 1200 PFU of virus or medium-only controls were added to the apical surface to a final volume of 100 μL. Viral supernatant was removed after 1 hour incubation at 37°C and infection was followed for the indicated time points. Viral production was then quantified by qRT-PCR using 3 consecutively collected apical washes of 100 μL PBS.

### Quantification of SARS-CoV-2 RNA by qRT-PCR

For quantification of viral RNA by qRT-PCR, total RNA was isolated using the High Pure Viral RNA kit (Roche) according to the manufacturer’s instruction. Viral RNA was quantified using GoTaq® 1-Step RT-qPCR kit (Promega). SARS-CoV-2 N gene RNA was amplified using forward (Ngene F cgcaacagttcaagaaattc 28844-28864) and reverse primers (Ngene R ccagacattttgctctcaagc 28960-28981). Copy numbers were calculated from a standard curve produce with serial 10-fold dilutions of SARS-CoV-2-RNA. Amplification program began with the RT-step 15 min at 50°C then the denaturation step 10 min at 95°C, and 10 s at 95°C, 10 s at 60°C and 10 s at 72°C (40 cycles). The melting curve was obtained by temperature increment 0,5°C/s from 60°C to 95°C.

### Interferon ELISA

Human IL-29/IL-28B (IFN-lambda 1/3) concentration in SARS-CoV-2 infected epithelium basal media was quantified using ELISA technics following manufacturer’s recommendations (R&D systems, Minneapolis, USA). 100 μl of media was used for each point.

### Immunofluorescence detection, antibodies and confocal microscopy

For antigen detection, BE were washed repeatedly with PBS to remove mucus then fixed with 4% paraformaldehyde for 30min using complete insert immersion. Epithelia were then washed and permeabilized with 0.5% TritonX-100 in PBS for 10min at room temperature and blocked in IF buffer (PBS containing 10% SVF and 0.5% saponin) for 1h at room temperature. Primary antibody and fluorescently labeled phalloidin to stain the actin cytoskeleton was diluted in IF buffer and applied to inserts for 1h at room temperature. Samples were washed three times under agitation with PBS and incubated with secondary antibody diluted in IF buffer and incubated for 2h at room temperature. Insert were then washed in PBS, desalted in H_2_O miliQ and rinsed in Ethanol 100% and air-dried. Membranes were then removed from inserts and mounted in DAPI (4’,6-diamidino-2-phenylindole) containing DAKO Fluorescence Mounting Medium prior to microscopy analysis. Mounted samples were subsequently examined on an epifluorescence microscope (Leica inverted DRMi6000 widefield microscope) at low resolution for kinetic studies. High resolution analysis was performed on a SP8 confocal microscope (Leica Microsystems at the Bordeaux Imagery center) using maximal pixel resolution at 20x, 40x or 63x respectively and 0.3μm Z-stacks resolution. Full epithelia overviews were acquired with Leica LAS-X software in spiral mosaics mode and three-dimensional reconstructions were done with Leica LAS-X software in 3D-viewer mode. Image processing was done using Image J software. Signal of interest were quantified using a semi-automatic macro. Briefly, Z-projections of different focal planes were generated and regions of interest (ROI) were manually inserted. Signal of interest was quantified automatically in each ROI, with appropriate predefined threshold and sizing for each condition. Quantification were performed to measured either number or size of signal of interest. Obtained values are represented either as absolute number or as normalized values (as indicated). The following primary antibodies and IF dilutions were used in this study; mouse monoclonal Ab anti-SARS-CoV-2-N clone 3G9 (this study, 1:500), rabbit monoclonal Ab anti-human Cytokeratin 5 (Abcam, ab52635, 1:200), rabbit polyclonal Ab anti-human Acetylated tubulin (Cell Signaling, D20G3, 1:200), rabbit polyclonal Ab anti-human ACE2 (Abcam, ab15348, 1:50), rabbit monoclonal Ab antihuman-Mucin 5AC (Abcam, ab198294, 1:200). The following secondary antibodies were used in this study; cross absorbed Donkey anti-mouse Alexa Fluor 488 or 647 (Life technologies, A212020/A31571, 1:300) and cross absorbed Donkey anti-rabbit Alexa Fluor 594 (Life technologies, A31573, 1:300) as well as Alexa-Fluor 594 labeled phalloidin (Invitrogen, 1:500).

### Electron microscopy

For electron microscopy, epitheliums were first washed in physiological serum and then fixed with 2.5% (v/v) glutaraldehyde and 2% (v/v) paraformaldehyde in 0.1M phosphate buffer (pH 7.4) during 2h minimum at room temperature (RT). Then samples were washed in 0.1M phosphate buffer and post-fixed in 1% (v/v) osmium tetroxide in phosphate buffer 0.1 M during 2h, in the dark, at RT, then washing in water and dehydrated through a series of graded ethanol and embedded in a mixture of pure ethanol and epoxy resin (Epon 812; Delta Microscopy, Toulouse, France) 50/50 (v/v) during 2 hours and then in 100% resin overnight at RT. The polymerization of the resin was carried out over a period between 24-48 hours at 60°C. Samples were then sectioned using a diamond knife (Diatome, Biel-Bienne, Switzerland) on an ultramicrotome (EM UCT, Leica Microsystems, Vienna, Austria). Ultrathin sections (70 nm) were picked up on copper grids and then stained with uranyless and lead citrate. Grids were examined with a Transmission Electron Microscope (H7650, Hitachi, Tokyo, Japan) at 80kV.

## Acknowledgments

We thank the staffs of both the pathology and surgery departments (both from the University Hospital of Bordeaux), Isabelle Goasdoue, Virginie Niel, and Marine Servat from the clinical investigation center, for technical assistance. Light and electron imaging was performed on the Bordeaux Imaging Center, member of the FranceBioImaging national infrastructure (ANR-10-INBS-04). We thank the BIC relentless support during the lockdown. We thank the “Agence Nationale de la Recherche” (ANR, ROSAE project CE14-0015-01) and ANR and the Fondation de France (ANACONDA project) for funding support. D.R.R is funded by the LabEx ParaFrap (ANR-11-LABX-0024). ML Blondot was supported by the Region Nouvelle Aquitaine and UBReact Bordeaux University. N. Pied is supported by the FRM DEQ20180339229. We thank the CNRS and the Bordeaux University for support. We are indebted to the members of our research teams, our families and several colleagues for their resilience and support during the past year. H. Wodrich is an INSERM fellow.

---

**Figure S1.**
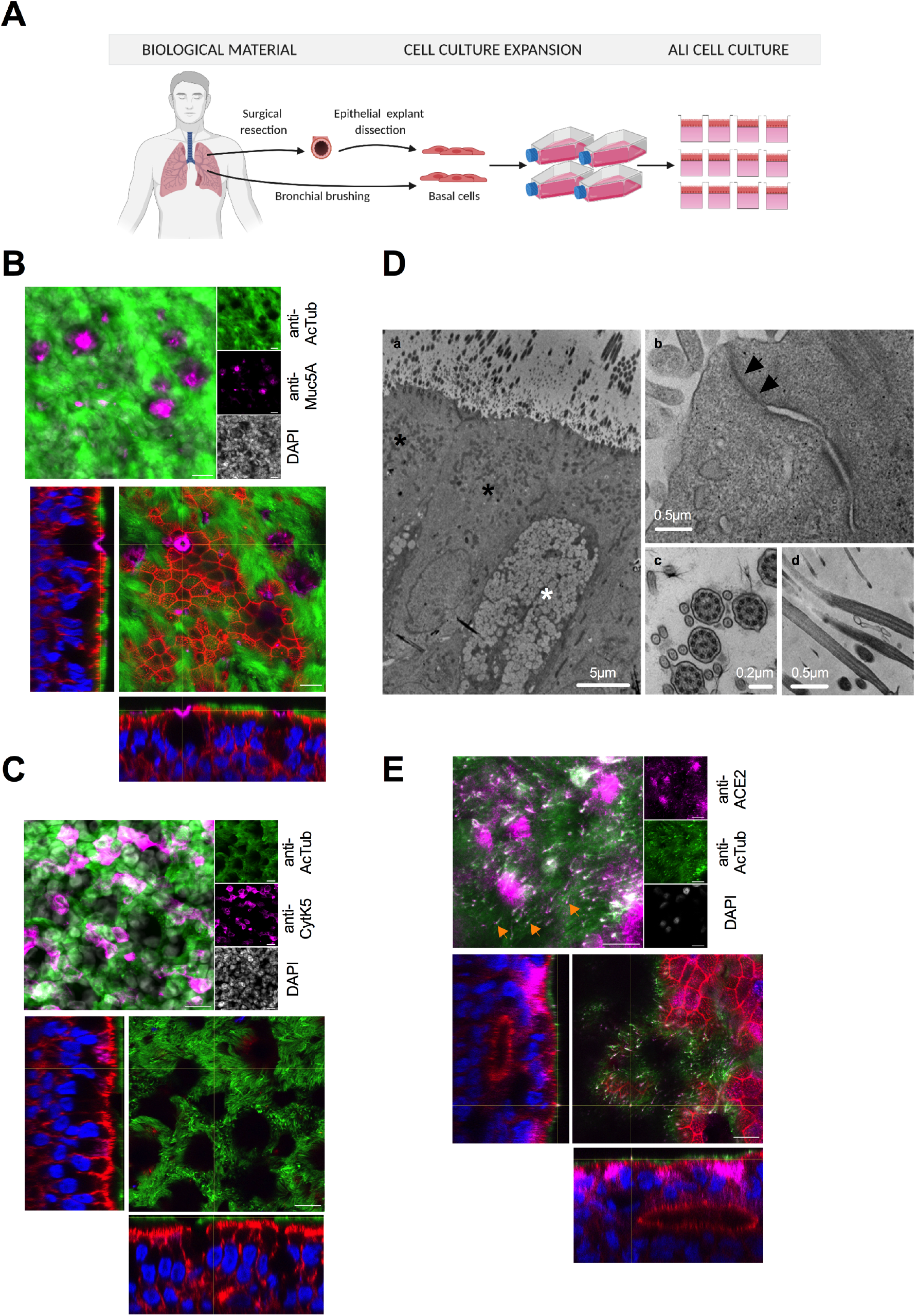
Characterization of bronchial epithelia (BE). A: Schematic overview of BE generation. Basal cells extracted from surgical dissection or bronchial brushing were expanded and differentiated at the air-liquid interface. B: Differentiated BE were stained with anti-acetylated tubulin to identify ciliated epithelia cells (green signal) or anti-Muc5A to detect cells (pink signal) and counterstained with DAPI (grey in top image, blue in bottom image). Top image shows a Z-projection, the bottom image shows an individual Z-section of a 3D reconstruction counterstained with phalloidin to detect the cell morphology via the actin cell cortex (red signal). Scale bar is 10μm. Note that ciliated cells are located to the apical side (see movie S1 for 3D). C: As in B but the differentiated BE was stained with with anti-acetylated tubulin (green signal) or anti-cytokeratin 5 to detect basal cells (pink signal) and counterstained with DAPI. Scale bar is 10μm. (see movie S2 for 3D). D: Electron microscopy of fully differentiated BE. The overview (a) shows ciliated epithelia cells (black asterisk) and goblet cells (white asterisk). The magnified images show tight junctions (b) marked by arrows and cilia either as cross-section (c) or longitudinal section (d). Scale bars are indicated. E: As in B but the differentiated BE was stained with with anti-acetylated tubulin (green signal) or anti-ACE2 to detect the SARS-CoV-2 receptor (pink signal) and counterstained with DAPI. Note that arrows point at individual cilia with ACE2 signal. Scale bar is 10μm. (see movie S2 for 3D).

**Figure S2.**
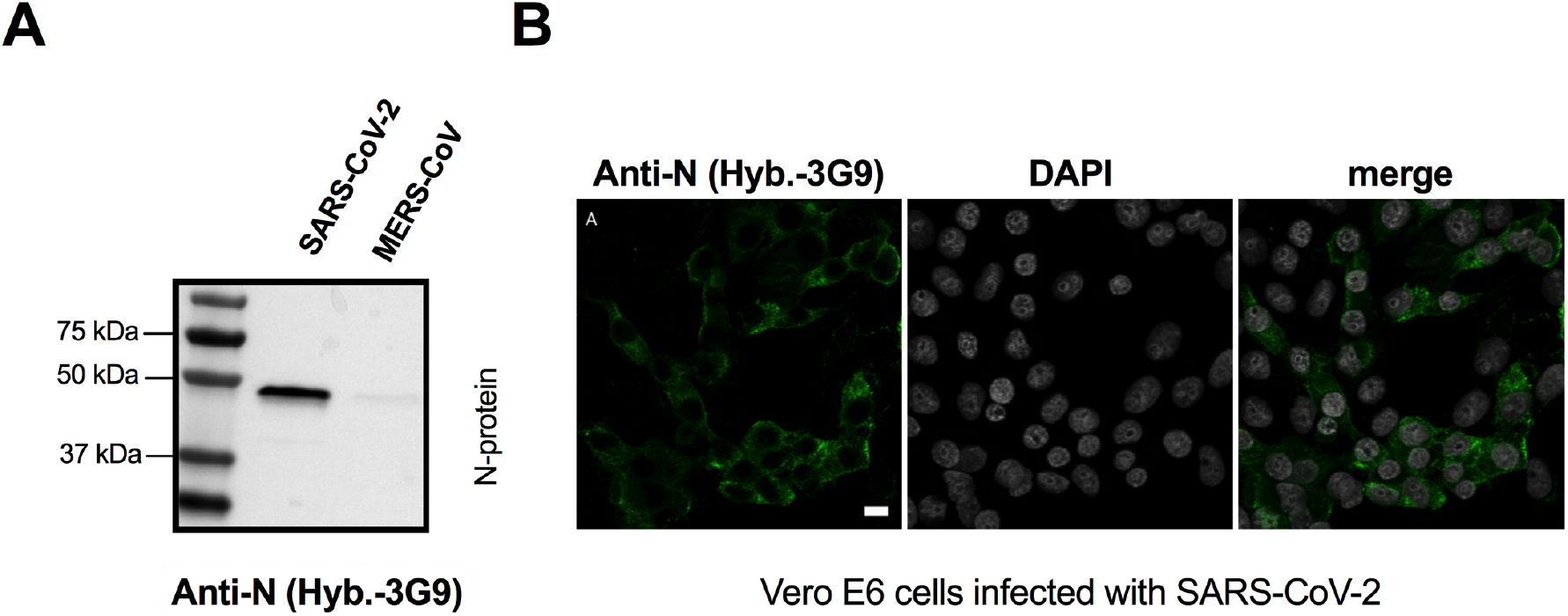
Characterization of monoclonal anti-SARS-CoV-2-N antibody (clone 3G9). A: Western blot analysis of recombinant bacterially purified SARS-CoV-2-N (100 ng, left lane) vs. MERS-CoV-N (100 ng, right lane). B: Detection of infected Vero E6 cells. Cells were infected for 24h with SARS-CoV-2, fixed and stained with monoclonal antibody to the nucleoprotein of SARS-CoV-2 (hybridoma 3G9).

**Figure S3.**
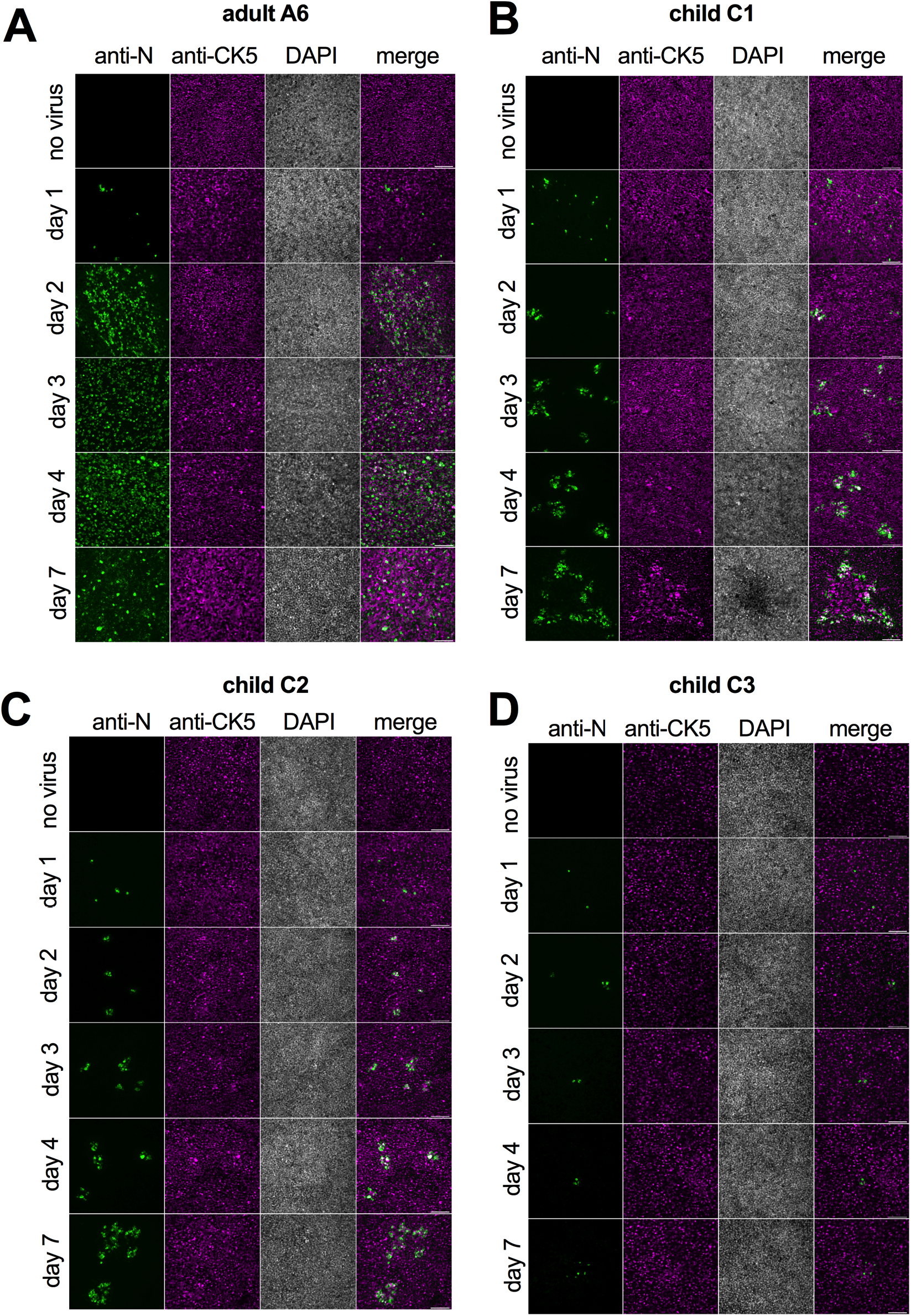
SARS-CoV-2 infection kinetic of bronchial epithelia (BE) from children and adult donor. A: Representative widefield microscopy images of BE from one adult donor and three children (A6 top left, C1 top right, C2 bottom left and C3 bottom right) at low resolution. BE were fixed at day 1, 2, 3, 4, 7 as indicated to the left of each row, non-infected controls were also fixed at day 7. BE were stained with anti-N antibodies to detect infected cells (green signal first column), anti-cytokeratin 5 to detect basal cells (magenta signal, second column) and counterstained with DAPI (grey signal, third column) and a merge of the three signals (forth column). Note the slow virus spread in the children derived epithelia. Scale bar is 10μm

